# Lifetime physical activity and network attack tolerance contribute to the preservation of motor function in Parkinson’s disease

**DOI:** 10.1101/2025.03.14.643226

**Authors:** Adrian L. Asendorf, Elena Guerra, Verena Dzialas, Magdalena Banwinkler, Hendrik Theis, Niklas Hagemann, Kathrin Möllenhoff, Thilo van Eimeren, Merle C. Hoenig

## Abstract

We tested whether network resilience, quantified by network attack tolerance (NAT), is associated with dopamine terminal (DaT) integrity, motor function and lifetime factors in Parkinson’s disease (PD). Data from 22 PD patients and 39 healthy controls included information on lifetime physical activity (PA), cognitive/motor performance, putaminal DaT integrity, and resting-state fMRI. NAT was assessed at global and subnetwork level by calculating global efficiency upon iterative node removal. Generalized linear-mixed-effects models were used to test the effects of PA, education, and dopamine integrity on NAT. Next, the moderating effect of lifetime factors on the association between NAT and motor function were assessed, controlling for DaT integrity. Greater putaminal DaT integrity was linked to higher somatomotor NAT. Higher global and somatomotor NAT supported motor function, especially in patients with moderate lifetime PA. Lifestyle factors may thus serve network-specific attack tolerance, thereby promoting motor preservation in PD, independent of dopaminergic impairment.

## Introduction

In Parkinson’s disease (PD), the severity of cognitive and motor symptoms can vary significantly between individuals, despite similar levels of dopaminergic impairment. This points towards mechanisms that partially mitigate the effects of progressive loss of dopaminergic innervation and regional neurodegeneration on functional performance. These compensatory or coping mechanisms have been summarized under the umbrella term of *resilience* and can be facilitated by lifestyle factors such as lifetime physical activity (PA) or education. By definition, resilience can be divided into the concepts of motor reserve (MR) and cognitive reserve (CR). While MR is linked to the preservation of motor function, for instance, by the adaptation of motor-relevant networks, CR is associated with the relative maintenance of cognitive performance through the adaptation of cognitive-relevant networks ^1,2^. To date, the neurobiological underpinnings of MR and CR in PD remain largely unknown.

Emerging evidence suggests improved efficiency, flexibility, and integration of functional networks to be associated with resilience in PD ^3–6^. Notably, a metric that has lately been proposed to reflect CR in PD is network attack tolerance (NAT) ^7^. NAT describes the ability of a functional network to maintain its topological integrity even when parts of the network are artificially removed ^8^. In graph theory, the brain is conceptualized as a network of nodes (regions) connected by edges (functional connections). NAT is assessed by systematically deleting nodes (i.e., attacks) from this network and evaluating whether the network sustains efficient communication. Concomitantly, networks that maintain their efficiency despite successive attacks represent high NAT. How efficiently information is exchanged in a network can be estimated by global efficiency (GE), which is assesses the shortest path length between all pairs of nodes in a network ^9,10^. Importantly, unlike other graph theoretical metrics, GE can account for paths of disconnected nodes, rendering it especially suitable for disconnected networks ^8,11^ and hence, a robust measure of efficiency for NAT.

In the context of PD, NAT may provide a valuable framework to study the brain’s actual compensatory capacity towards a decreasing dopaminergic input and progressive neurodegeneration. Indeed, higher NAT in the frontoparietal network was recently associated with the absence of cognitive decline in PD patients ^7^. However, it remains unknown whether NAT of certain brain networks is directly impacted by dopamine loss in PD. Furthermore, given that the cardinal symptoms of PD relate to motor impairment, it remains to be investigated whether greater NAT is also associated with the preservation of motor function (i.e., MR).

Notably, individual differences in resilience capacity, such as in MR or CR, have consistently been associated with certain lifestyle factors, such as education and PA. In PD, education has been positively associated with CR ^12–14^, while PA has traditionally been implicated as a proxy for MR. Both intense exercise interventions ^15,16^ and higher lifetime/habitual PA ^17,18^ have been demonstrated to attenuate motor decline and enhance dopamine availability ^18,19^ (for a recent review see ^20^). However, whether these lifestyle factors (i.e., reserve proxies) contribute to greater NAT remains unknown.

In this study, we therefore aimed to address three objectives: (1) to examine the impact of putaminal dopamine transporter (DaT) integrity on global and subnetwork NAT; (2) to directly test the association between NAT and two lifestyle factors (i.e., PA and education); while accounting for DaT integrity; and (3) to subsequently investigate the moderating effects of the lifestyle factors on the association between NAT and general motor and cognitive performance. These objectives were tested in a cohort of early PD patients, who were part of the “Dopamine and Motor Control” (DoMoCo) study that encompasses an extensive array of lifestyle questionnaires, DaT single-photon emission computed tomography (SPECT), MRI imaging, and a detailed battery of cognitive and motor function tests. We hypothesized that preserved integrity of the dopaminergic system has a positive effect on NAT, particularly in the somatomotor network. We further hypothesized that, despite differences in dopaminergic impairment, lifetime PA (MR proxy) would be linked to greater NAT in the somatomotor network, whereas education (CR proxy) would be associated with greater NAT in cognition-relevant networks, such as the frontoparietal or attention network. Finally, we hypothesized that lifetime PA and education would moderate the association between NAT and general motor and cognitive performance, respectively.

## Materials and Methods

### Participants

This study comprised 22 patients with early PD, who were recruited for the DoMoCo study during their clinical workup at the Department of Nuclear Medicine at the University Hospital Cologne, Germany. The DoMoCo study is conducted as part of the Collaborative Research Centre 1451 (CRC 1451) at the University Hospital Cologne. The inclusion criteria for this study were: 1) age between 49 and 80 years; 2) early-stage PD (clinical symptoms <3 years); 3) available in-house DaT SPECT imaging; 4) right-handedness; 5) absence of severe cognitive deficits (Montreal Cognitive Assessment (MoCA) score >= 24); 6) absence of depressive symptoms (Geriatric Depression Scale score <= 5); 7) no current administration of selective serotonin reuptake inhibitors; 8) absence of significant other neurological disorders; 9) MRI safe (e.g. no pacemaker). Notably, all tests and imaging procedures in the DoMoCo study were performed in PD patients off medication. As 13 of the 22 patients had already received dopaminergic medication, they were asked to discontinue their medication prior to study participation. This represented at least 12 hours of withdrawal of dopamine replacement therapy or 72 hours for extended release of dopamine agonists.

A group of 39 healthy controls, who underwent the study protocol of the DoMoCo study, except the DaT SPECT imaging, was used as reference for the cognitive and motor tests.

The study was approved by the local ethics committee (reference: 20-1422) and was conducted according to the standards of the Declaration of Helsinki. All participants gave informed consent prior to enrollment and received monetary compensation for their participation. Demographic characteristics are summarized in Table 1.

**Table 1:** Demographics of the study cohort. Standard deviation (SD) is provided for continuous variables and interquartile range (IQR) for ordinal values (MDS UPDRS-III and MoCA scores). While the z-scores for DaT integrity were derived from an internal software cohort, the z-scores for GCP and GMP were based on the 39 HCs described in this table. PD = Parkinson’s disease, HC = healthy control, DaT = dopamine transporter, PA = lifetime physical activity, MoCA = Montreal Cognitive Assessment, GCP = general cognitive performance, GMP = general motor performance, MDS UPDRS-III = Movement Disorder Society Unified Parkinson’s Disease Rating Scale-Part III, LEDD = Levodopa equivalent daily dose.

|  | Mean (SD) /Median (IQR) |  |
| --- | --- | --- |
|  | PD (n = 22) | HC (n= 39) |
| Age | 62.1 (8.0) | 63.5 (6.5) |
| Sex (m/f) | 15/7 m =68% | 26/13 m= 50% |
| Putaminal DaT integrity (z-scores) | -3.6 (0.7) | NA |
| Education (years) | 15.8 (2.7) | 16.7 (2.8) |
| PA | 58.3 (82.8) | 48.1 (34.0) |
| MoCA | 27 (2) | 28 (2) |
| GCP (z-scores) | -0.4 (2.4) | NA |
| GMP (z-scores) | -3.2 (3.1) | NA |
| MDS UPDRS III | 14 (6) | 0 (1) |
| LEDD | 206.4 (113.2) | NA |

### Reserve proxies – Lifetime physical activity and education

Lifetime PA was used as a proxy for MR and assessed using the Historical Leisure Activity Questionnaire (HLAQ) ^21^ and metabolic equivalents (METs) ^22^, which were assigned to account for the intensity of the respective activity. The HLAQ consists of four periods (12-19 years, 19-35 years, 35-50 years, above 50 years). For each period, the participant had to indicate which activity he or she performed more than 10 times and specify the number of hours per week, months per year, and years per period. Using the formula below, we first calculated the average weekly number of hours each activity was carried out per period. Next, we obtained the weekly energy expenditure for each activity by multiplying the average weekly hours per activity by the corresponding MET score. Finally, a total weekly energy expenditure score (WEE_period_) was calculated over all activities reported in that period. In this study, lifetime PA was defined as the mean of the WEE_period_ calculated for the periods 19-35 years and 35-50 years. We excluded the earliest period of the questionnaire (i.e., 12-19 years), given that a recall bias was noticed across participants. We also excluded the latest period (i.e., above 50 years) because all patients were only recently diagnosed with PD at around 50 years of age, which may have influenced their physical activity performance.

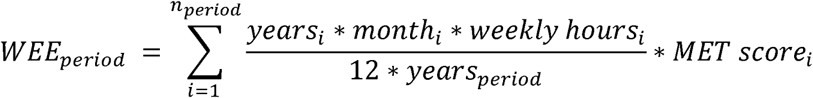

Aside from the lifetime PA variable, we further used educational attainment as proxy for CR. Educational attainment was defined as the total number of years spent in formal schooling, vocational training and/or university education.

### Behavioral assessment - Cognitive and motor function

All participants in the DoMoCo study underwent a detailed clinical, motor, and cognitive assessment. Clinically, disease severity was evaluated using the Movement Disorder Society Unified Parkinson’s Disease Rating Scale (MDS-UPDRS) Part III ^23^. To obtain more quantitative measures for cognitive and motor performance that go beyond clinical severity ratings, a battery of motor and cognitive tests was included in the current analysis. The test battery was conducted in the context of the Human Motor Assessment Centre (CRC 1451) and consisted of the following tests: Motor function was evaluated by measuring maximum grip strength and through the Finger Tapping Test, Purdue Pegboard Test, Jebsen-Taylor Hand Function Test, and Timed Up and Go Test. With exception of the Timed up and Go Test ^24^, a detailed description of the motor battery can be found elsewhere ^25^. The cognitive battery consisted of the Cologne Neuropsychological Screening Tool for Stroke ^26^, DemTect Test ^27^ and Trail Making Test ^28^. To obtain a measure for general motor performance (GMP) and general cognitive performance (GCP), the performance scores of the motor and the cognitive test battery were z-standardized using the sample of 39 age- and sex-matched healthy controls as reference. Subsequently, these z-scores were combined into two single composite scores, GMP and GCP, respectively. As longer durations in the Jebsen-Taylor, Timed Up and Go, and Trail Making Test represented poorer performance, these test scores were inverted prior to the composite score calculation.

### Dopamine transporter SPECT acquisition and preprocessing

DaT SPECT images were acquired on a PRISM-3000 three-head SPECT system (Philips/Picker) located at the University Hospital of Cologne following a standardized clinical procedure (i.e., [^123^I]Isoflurane injection, reconstruction using Chang’s attenuation correction, and voxel value normalization to the occipital cortex. Using the EARL-BRASS software (Hermes, Sweden; Tossici-Bolt et al., 2017), z-transformed deviation maps are provided that are based on a reference sample of age-matched healthy controls. Finally, for the current analysis, mean z-values of the bilateral putamen were extracted using EARL-BRASS.

### fMRI acquisition

All participants underwent a MRI session at the Research Centre Juelich. Anatomical T1- weighted MRI and resting state (rs) functional MRI (fMRI) series were acquired on a 3T Siemens Prisma scanner. The anatomical sequence followed a specific protocol: repetition time (TR) = 2300 ms, echo time (TE) = 2.32 ms, flip angle = 8°, slice thickness = 0.90 mm, voxel size = 0.9 x 0.9 x 0.9 mm, number of slices = 192. For each individual, one of three different fMRI sequences was available. Five individuals were scanned with the following settings: TR = 800 ms, TE = 37 ms, multiband SMS factor = 8, flip angle = 52°, slice thickness = 2 mm, voxel size = 2.0 x 2.0 x 2.0 mm, number of volumes 740, acquisition time = approx. 10 min. While 16 individuals were scanned with these settings: TR = 980 ms, TE = 30 ms, multiband SMS factor = 4, flip angle = 70°, slice thickness = 2.2 mm, voxel size = 2.2 x 2.2 x 2.2 mm, number of volumes 500, acquisition time = approx. 8 min. Finally, one individual was scanned with a sequence including: TR = 1120 ms, TE = 57 ms, multiband SMS factor = 8, flip angle = 52°, slice thickness = 2 mm, voxel size = 2.0 x 2.0 x 2.0 mm, number of volumes 740, acquisition time = approx. 13 min. The sequence changes were unintended, but necessary due to an unforeseen, minor update of the scanner’s software during the study period.

### fMRI preprocessing

Preprocessing of the fMRI data was carried out using the CONN toolbox v20.b (Whitfield-Gabrieli & Nieto-Castanon, 2012) implemented in Matlab (Matlab R2020b Update 4, MathWorks, Inc., Natick, Massachusetts, United States). CONN’s default preprocessing pipeline was used to normalize the images to MNI (Montreal Neurological Institute) space (web.conn-toolbox.org/fmri-methods/preprocessing-pipeline). The pipeline included spatial realignment, slice-timing correction, outlier identification, segmentation, normalization, and smoothing (Gaussian kernel 8mm full-width half maximum). The preprocessing was performed separately for each of the three sequences.

### Graph theoretical network construction and network attack

In a first step, we used a set of 300 predefined, spherical, non-overlapping regions of interest (ROIs) ^30^ to compute individual ROI-to-ROI connectivity matrices in CONN. These 300 x 300 covariance matrices represent the functional connectivity between each ROI pair. Each element or tile of these covariance matrices contained the respective Fisher-transformed bivariate correlation coefficient between the Blood Oxygen Level-Dependent (BOLD) time series of two ROIs.

In a second step, we computed weighted, undirected, and unthresholded graphs based on the covariance matrices utilizing the Brain Connectivity Toolbox (brain-connectivity-toolbox.net) ^8^ and the Graph Analysis Toolbox (cbrain.stanford.edu/Tools.html) ^31^. These graphs were computed on the whole-brain level as well as the single subnetwork level. The single subnetworks of interest were the default mode network (DMN, 65 ROIs), the frontoparietal network (FPN, 36 ROIs), the somatomotor network (SMN, 51 ROIs), and the attention network (ATN, 27 ROIs), which are the major large-scale cognitive and motor networks defined by various resting state studies ^32–34^. Subsequently, each of these graphs was proportionally thresholded, preserving only the strongest ten to fifty percent of the weighted edges in the graph, respectively. We chose this broad range as there is no consensus on appropriate thresholds for NAT analyses. Consequently, by using five percent increments, nine different network densities per network were examined. Each of these graphs was binarized, leaving nine differently thresholded, unweighted, undirected graphs per network and individual. Figure 1 provides a graphic outline of the analysis and Supplementary Figure 1 portrays the approach as a detailed procedural map.

**Figure 1:**
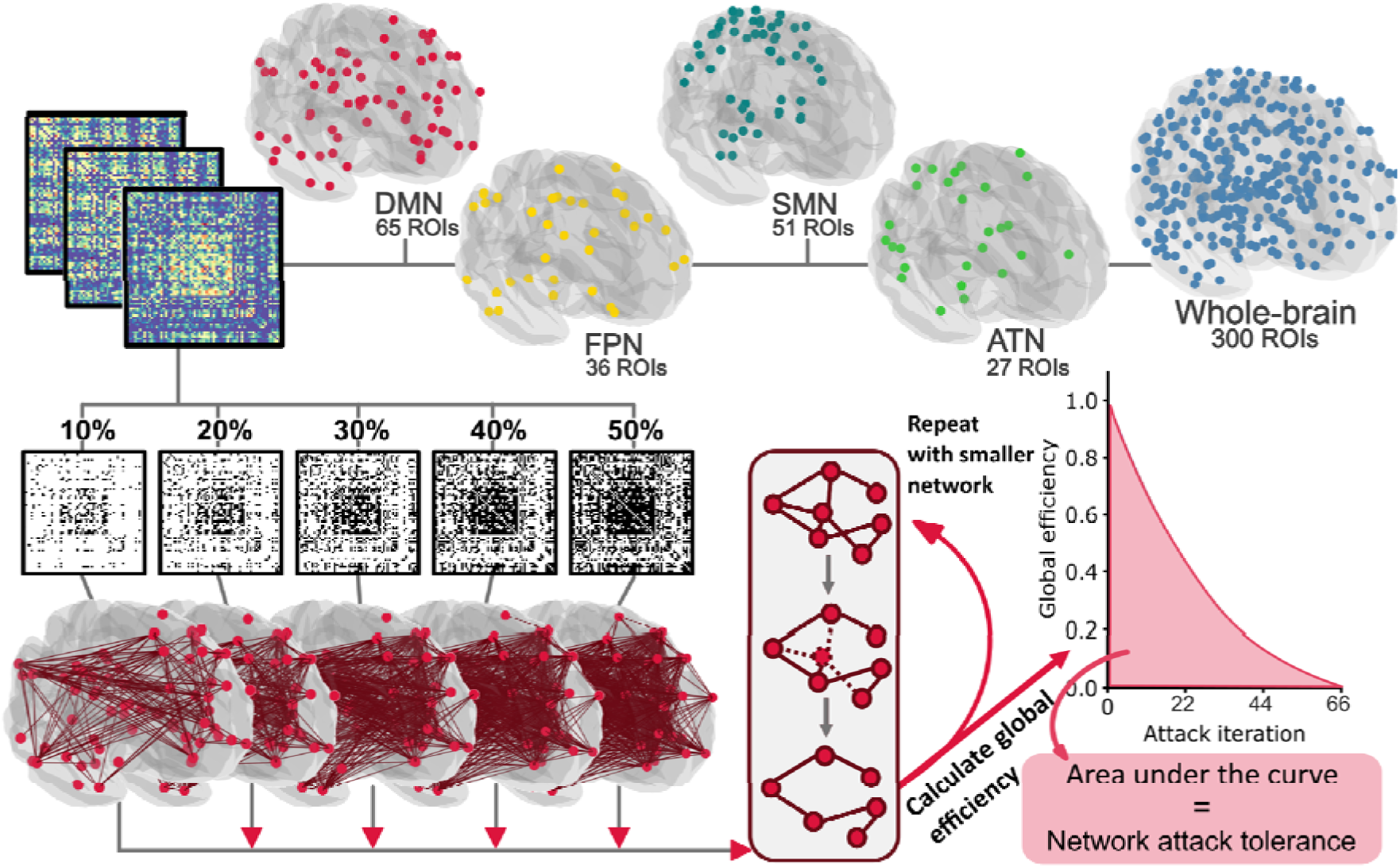
Illustration of the analysis pipeline: **Top:** Individual resting-state ROI-to-ROI connectivity matrices were computed using ROIs based on a functional atlas. ROIs of the attention network (ATN, green), the default mode network (DMN, red), the frontoparietal network (FPN, yellow), the somatomotor network (SMN, turquoise) and the whole-brain network (multiple colors) were analyzed separately. The analysis steps are illustrated using the DMN as an example. **Bottom left:** Connectivity matrices were thresholded at nine different densities from 10% to 50% (for clarity only five out of nine densities are depicted above), then binarized, and finally used for network construction. **Bottom right:** During the attack analysis, the network’s nodes were iteratively removed in descending order of their connectedness. For each removed node, the network’s global efficiency was computed. The ability to sustain its global efficiency to successive attacks (area under the curve) defined the attack tolerance of the respective network. All renderings and covariance matrices were plotted using nilearn ^35^

In a final step, the constructed graphs were attacked using the same set of toolboxes. During the attack analysis, nodes were iteratively removed from the network in descending order of their degree (i.e., the number of edges connected to that node). Thus, nodes with the most connections were removed first. After each node removal, the network’s GE was calculated. GE was defined by the average inverse shortest path length between all pairs of nodes in the network, where N represented the number of nodes in the network and L_ij_ was the shortest path between nodes i and j ^9^:

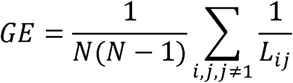

GE was then plotted on a graph for each attack iteration (see Figure 1). The area under the curve (AUC) of each graph was used to obtain one single measure of individual network attack tolerance (NAT). Next, the AUC was normalized according to the number of attacks performed (i.e., the number of nodes in the respective network). After normalization, NAT described how many percent of the maximum possible GE was retained during attacks.

### Code availability

The underlying inhouse code for this analysis is available on GitHub and can be accessed via this link https://github.com/Asendorfa/NAT. We provide a procedural map in the supplements that depicts which script was used for which analysis step (Supplementary Fig. 1).

## Statistical analyses

### Generalized linear mixed-effects model

In a first step, we wanted to examine the direct effects of two reserve proxies, specifically lifetime PA and education, as well as the effects of dopaminergic impairment, namely putaminal DaT integrity, on NAT, while accounting for individual variances in demographics. To account for the repeated measure design due to the different network densities, we performed generalized linear mixed-effects models (GLMMs) ^36^, allowing individual intercepts for each participant. Fixed effects included lifetime PA, education, and putaminal DaT-integrity, while controlling for network density, fMRI sequence (interscan variability), age, and sex. Since NAT is a fraction, a beta regression model ^37^, i.e., a GLMM with a beta response distribution, utilizing a logit link function was used. R-squared estimates of the GLMMs were derived from the ‘mgcv’ R package ^38^.

### Moderation analysis

Next, we assessed the potential moderating effects (thus indirect effects) of lifetime PA and education on the association between NAT and motor/cognitive performance. To do this, we ran two linear moderation models estimating either general motor or cognitive performance based on the respective network’s NAT, the reserve proxies (education and PA) as well as their interaction. Since NAT was originally a repeated measurement (due to the nine different network densities), we used the mean NAT value across all densities for each network as input variable. More specifically, in the first model we estimated motor performance (GMP) using putaminal DaT integrity, age, sex, fMRI sequence, education, and NAT, introducing PA as moderating variable. The second model estimated cognitive function (GCP) using putamen DaT integrity, age, sex, fMRI sequence, PA, and NAT as covariates and education as a moderating variable. Since GCP was missing for one individual this model was run on data of 21 individuals. Finally, the Johnson-Neyman technique ^39^ was used to identify ranges of significance and visualize interaction effects in the data.

All statistical models listed above were run separately for global and subnetwork NAT (5 runs) using R (Rstudio ver. 2022.12.0) and the following packages: ‘mgcv’ ^38^, ‘ggplot2’ ^40^, ‘interactions’ ^41^, ‘broom’ ^42^.

## Results

### Effect of DaT integrity and reserve proxies on NAT

The GLMMs assessing the contributions of DaT integrity and reserve proxies on NAT, while controlling for demographic, network density and interscan variability, yielded a positive effect of putaminal DaT integrity (β=.079; *p*<.001) on SMN NAT (see Figure 2). In addition, one of our reserve proxies, namely education, presented a positive effect (β=.013; *p*=.006) on NAT in the ATN (i.e., higher education was associated with increased NAT in the ATN). Remaining models did not result in significant findings in terms of DaT integrity or reserve proxies.

**Figure 2:**
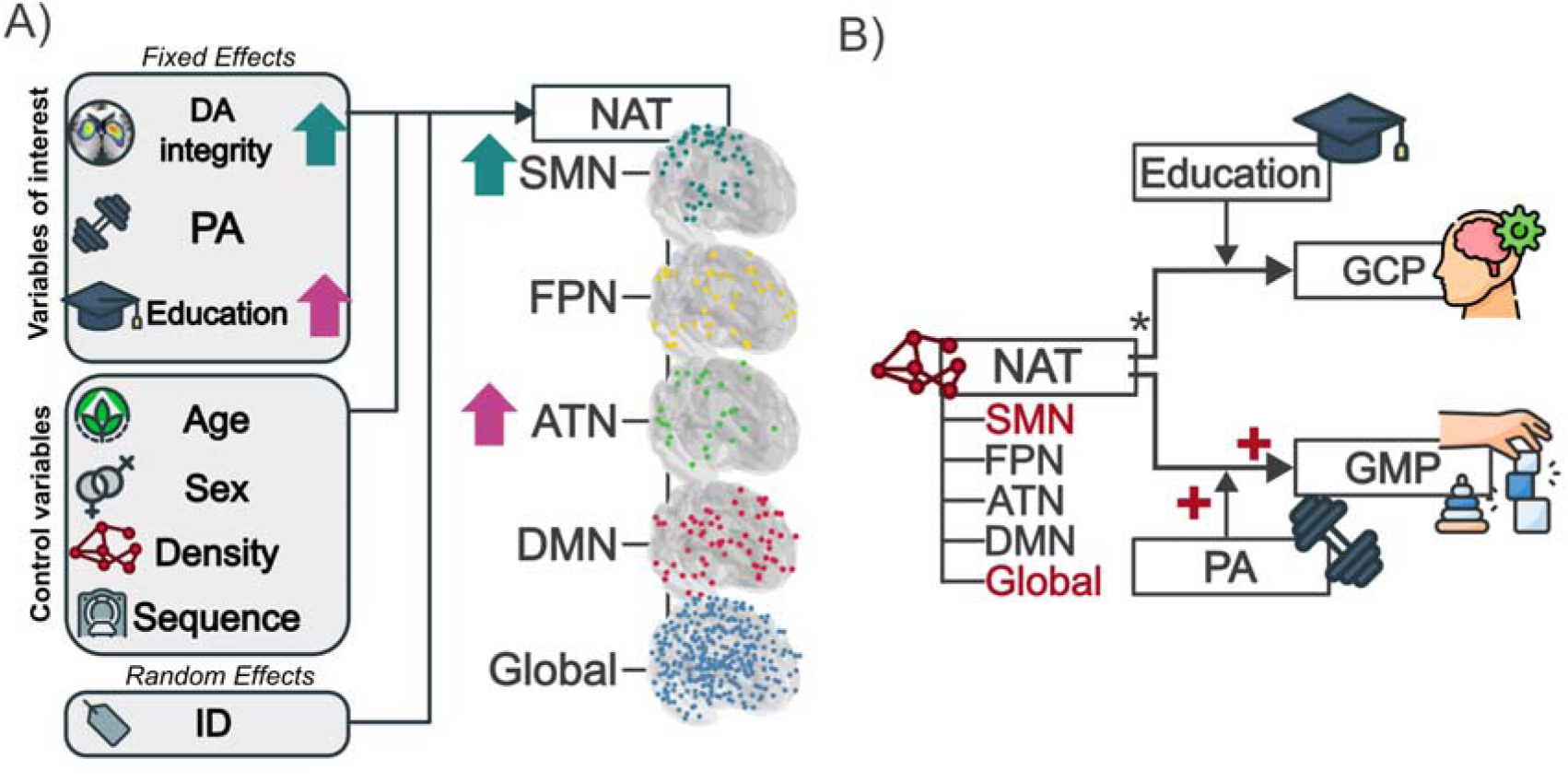
Model structure and key results. **A):** Results of the generalized linear mixed models estimating the effect of biological (putamen dopamine terminal (DaT) integrity) and reserve factors (education and physical activity (PA)) on NAT. The arrows are color-coded and represent the direction of the association between NAT of the respective network and the covariate. The attention network (ATN) is shown in green, and the somatomotor network (SMN) in turquoise. **B)**: Results of the regression models. *Top:* Regression model predicting general cognitive performance (GCP) using education as a moderator. *Bottom:* Regression model predicting general motor performance (GMP) using PA as a moderator. The plus sign indicates a positive effect exerted by NAT (corresponding network also in red) and PA as a moderator on GMP. *Control variables for moderation analyses: age, sex, putaminal DaT, education or PA. For better visualization, effects of the control variables are not shown in both illustrations.

Regarding the control variables, we found a positive effect of age (β=.001; *p*=.024), network density (β=.128; *p*<.001) and interscan variability (1<2: β=.035; *p*=.020, 1<3: β=.059; *p*=.026) on global NAT. For the SMN, there was a positive effect of density (β=.593; *p*<.001) and interscan variability (1<2: β=.109; *p*=.005) on NAT. In terms of DMN NAT, a positive effect of sex (f<m: β=.071; *p=*.043) and network density (β=.565; *p*<.001) was observed. Regarding ATN NAT, the model yielded a negative effect of sex (f<m. β=.071; *p=*.043) and a positive effect of network density. Finally, FPN NAT was positively affected by network density (β=.671; *p<*.001) and interscan variability (1<2: β=.077; *p*=.014, 1<3: β=.198; *p*<.001). All results have been summarized in Table 2.

**Table 2:** Results table of GLMMs assessing the factors contributing to network attack tolerance (NAT) in five different networks. Significant findings are highlighted in bold. ß = unstandardized beta coefficients, *: p<.05, CI represents the 95% confidence interval, ATN = attention network, DMN = default mode network, FPN = frontoparietal network, SMN =somatomotor network

| Fixed effects | Global |  |  |  | SMN |  |  |  | DMN |  |  |  | ATN |  |  |  | FPN |  |  |  |
| --- | --- | --- | --- | --- | --- | --- | --- | --- | --- | --- | --- | --- | --- | --- | --- | --- | --- | --- | --- | --- |
| | $\beta$ | $p$ | CI | | $\beta$ | $p$ | CI | | $\beta$ | $p$ | CI | | $\beta$ | $p$ | CI | | $\beta$ | $p$ | CI | |
| Biological factor |  |  |  |  |  |  |  |  |  |  |  |  |  |  |  |  |  |  |  |  |
| Mean putamen | .008 | .289 | -.007 | .023 | **.079** | **<.001\*** | **.040** | **.118** | -.030 | .144 | -.069 | .010 | -.010 | .531 | -.042 | .022 | .014 | .395 | -.018 | .046 |
| Reserve proxies |  |  |  |  |  |  |  |  |  |  |  |  |  |  |  |  |  |  |  |  |
| Physical activity | .000 | .968 | .000 | .000 | .000 | .823 | .000 | .000 | .000 | .419 | .000 | .000 | .000 | .641 | .000 | .000 | .000 | .137 | .000 | .000 |
| Education | -.001 | .503 | -.006 | .003 | -.009 | .140 | -.020 | .003 | .004 | .444 | -.007 | .016 | **.013** | **.006\*** | **.004** | **.022** | .001 | .815 | -.008 | .010 |
| Control variables |  |  |  |  |  |  |  |  |  |  |  |  |  |  |  |  |  |  |  |  |
| Age | **.001** | **.024\*** | **.000** | **.003** | -.003 | .165 | -.007 | .001 | -.001 | .738 | -.005 | .003 | .001 | .686 | -.003 | .004 | .003 | .113 | -.001 | .006 |
| Sex(f<m) | .024 | .066 | -.002 | .051 | -.003 | .528 | -.090 | .046 | **.071** | **.043\*** | **-.002** | **.140** | **-.069** | **.015** | **-.125** | **-.014** | -.025 | .373 | -.080 | .030 |
| Network density | **.128** | **<.001\*** | **.119** | **.138** | **.594** | **<.001\*** | **.570** | **.618** | **.565** | **<.001\*** | **.544** | **.586** | **.754** | **<.001\*** | **.722** | **.786** | **.671** | **<.001\*** | **.645** | **.697** |
| Sequence (1<2) | **.035** | **.020\*** | **.005** | **.064** | **.109** | **.005\*** | **.033** | **.185** | .076 | .051 | .000 | .153 | -.008 | .795 | -.070 | .054 | **.077** | **.014\*** | **.016** | **.139** |
| Sequence (1<3) | **.059** | **.026\*** | **.007** | **.110** | .072 | .295 | -.063 | .207 | .087 | .209 | -.049 | .223 | .064 | .252 | -.046 | .174 | **.198** | **<.001\*** | **.090** | **.306** |
| Explained var. | R<sup>2</sup>=.81 N=198 |  |  |  | R<sup>2</sup>=.93 N=198 |  |  |  | R<sup>2</sup>=.94 N=198 |  |  |  | R<sup>2</sup>=.93 N=198 |  |  |  | R<sup>2</sup>=.94 N=198 |  |  |  |

### Effect of NAT on motor function and cognitive function

Given that we observed an effect of DaT integrity and one reserve proxy, we next aimed to test whether global and subnetwork NAT has an impact on behavior, namely motor performance (GMP) and cognitive performance (GCP), while considering a moderating effect of our respective reserve proxy (PA and education). The results are described below.

#### Effect of NAT on motor function

In terms of GMP, the linear moderation models yielded GMP to be positively affected by higher global NAT (β=279.127; *p*=.005) and SMN NAT (β=117.415; *p*=.017), respectively. Remaining models did not yield a significant effect of subnetwork NAT on GMP. Regarding the moderation effect of PA on this association, we found a trend significant result for SMN NAT (β=-1.418; *p=*.087). We further elucidated the moderation effects of PA on GMP using the Johnson and Neymann technique. The test showed a significant moderating effect of PA, which was only present at PA levels below 46.66 WEE for global NAT and 29.35 WEE for SMN NAT on GMP (see Supplementary Figure 2). Hence, at stationary NAT levels, PD patients with moderate PA levels demonstrated better GMP compared to highly sedentary PD patients suggesting that a moderately active lifestyle, in contrast to a more sedentary one, may enhance the positive impact of NAT on motor performance (see Figure 2B).

In terms of our covariates, we observed a significant effect of education on GMP in the ATN model (β=-.530; *p*=.033). Across the five tested moderation analyses, a negative effect of age on GMP was found (Global: β=-.348; *p*<.001; SMN: β=-.253; *p*=.002; DMN: β=-.288; *p*=.002, ATN: β=-.323; *p*<.001, FPN: β=-.294; *p*=.003). The model outputs are summarized in Table 3.

**Table 3:** Results table of models assessing the interaction of network attack tolerance (NAT) and physical activity (PA) on motor performance (GMP) in five different networks. (Trend) significant findings are highlighted in bold. ß = unstandardized beta coefficients, *: p<.05, CI represents the confidence interval, ATN = attention network, DMN = default mode network, FPN = frontoparietal network, SMN =somatomotor network

| Effects on GMP | Global |  |  |  | SMN |  |  |  | DMN |  |  |  | ATN |  |  |  | FPN |  |  |  |
| --- | --- | --- | --- | --- | --- | --- | --- | --- | --- | --- | --- | --- | --- | --- | --- | --- | --- | --- | --- | --- |
| | $\beta$ | p | CI | | $\beta$ | p | CI | | $\beta$ | p | CI | | $\beta$ | q | CI | | $\beta$ | p | CI | |
| Direct effects |  |  |  |  |  |  |  |  |  |  |  |  |  |  |  |  |  |  |  |  |
| NAT | **279.127** | **.005\*** | **102.564** | **455.69** | **117.415** | **.017\*** | **25.415** | **209.414** | 43.665 | .417 | -69.486 | 156.816 | 137.851 | .078 | -18.204 | 293.907 | 68.816 | .369 | -91.791 | 229.423 |
| PA | .792 | .114 | -.219 | 1.802 | .345 | .083 | -.053 | .743 | -.017 | .901 | -0.3 | 0.267 | .284 | .200 | -.172 | .741 | .243 | .279 | -.224 | .709 |
| Indirect effects |  |  |  |  |  |  |  |  |  |  |  |  |  |  |  |  |  |  |  |  |
| NAT x PA | -2.536 | .116 | -5.793 | .721 | **-1.418** | **.087** | **-3.075** | **.238** | .105 | .864 | -1.197 | 1.406 | -1.335 | .208 | -3.521 | .851 | -1.066 | .287 | -3.153 | 1.02 |
| Control variables |  |  |  |  |  |  |  |  |  |  |  |  |  |  |  |  |  |  |  |  |
| Education | -.206 | .212 | -.545 | .134 | -.124 | .507 | -.517 | .27 | -.375 | .101 | -0.836 | 0.086 | **-.53** | **.033\*** | **-1.007** | **-.052** | -.265 | .230 | -.722 | .192 |
| Mean putamen | .115 | .851 | -1.194 | 1.424 | -.402 | .629 | -2.172 | 1.367 | .282 | .709 | -1.326 | 1.891 | .513 | .484 | -1.032 | 2.057 | .455 | .595 | -1.359 | 2.27 |
| **Age** | **-.343** | **<.001\*** | **-.481** | **-.205** | **-.253** | **.002\*** | **-.39** | **-.116** | **-.288** | **.002\*** | **-0.449** | **-0.127** | **-.323** | **.001\*** | **-0.477** | **-.17** | **-.294** | **.003\*** | **-.463** | **-.124** |
| Sex (f<m) | -.700 | .483 | -2.807 | 1.407 | -.335 | .773 | -2.812 | 2.142 | -.239 | .86 | -3.136 | 2.658 | 1.079 | .41 | -1.675 | 3.834 | .315 | .803 | -2.374 | 3.003 |
| Sequence (1<2) | .338 | .769 | -2.115 | 2.791 | .748 | .577 | -2.095 | 3.592 | .977 | .517 | -2.211 | 4.166 | 1.751 | .184 | -.953 | 4.455 | 1.996 | .238 | -1.502 | 5.493 |
| Sequence (1<3) | -.66 | .091 | -8.003 | .683 | -1.443 | .498 | -5.945 | 3.059 | -2.234 | .386 | -7.643 | 3.175 | -2.298 | .333 | -7.257 | 2.662 | -1.767 | .57 | -8.357 | 4.823 |
| Explained var. | $R^2=.85$ N=22 | | | | $R^2=.82$ N=22 | | | | $R^2=.74$ N=22 | | | | $R^2=.78$ N=22 | | | | $R^2=.74$ N=22 | | | |

#### Effect of NAT on cognitive function

Concerning the association between GCP and NAT, none of the five linear moderation models demonstrated overall significance of the model test statistic, preventing interpretation of the individual effects of the moderation, and controlling variables.

## Discussion

In this study, we applied a multimodal neuroimaging approach (DaT SPECT, fMRI) combined with a detailed characterization of lifestyle, cognitive, and motor performance in a group of early PD patients. The overarching aim was to elucidate the role of a graph theoretical metric, namely NAT, as a potential motor and cognitive reserve measure that is linked to distinct lifestyle features. In this context, we identified NAT of the SMN as one potential MR mechanism that is related to lifetime PA (MR proxy), DaT integrity, and the relative preservation of general motor performance. In particular, we found that lower putaminal DaT integrity is directly associated with decreased robustness of the SMN against successive perturbations/attacks. In turn, regardless of the underlying DaT integrity, higher NAT of the SMN and global NAT were associated with better general motor performance, highlighting NAT’s crucial role in the relative preservation of motor function (i.e., MR). Importantly, we observed that the beneficial effect of SMN NAT on motor performance was less pronounced in individuals with a relatively sedentary lifestyle (i.e., lower PA levels) compared to moderately active individuals. This underlines the facilitating role of lifetime PA to the build-up of MR mechanisms, i.e., the strengthening of global and SMN NAT. Regarding CR, our findings indicated a direct effect of education on the NAT of a cognition-relevant network, namely the ATN. Yet, no moderating effect of education on the association between ATN NAT and general cognitive or motor performance was observed. Taken together, the results indicate NAT as crucial resilience mechanism, which may compensate for the progressive loss of dopaminergic terminals in PD, as will be discussed in the following.

### The influence of dopaminergic innervation on network attack tolerance

Importantly, and in line with our hypothesis, diminished DaT integrity was associated with lower SMN NAT. This observation was solely confined to the SMN, which contained four ROIs associated with the basal ganglia. These results suggest that a diminished dopaminergic innervation of the basal ganglia may lead to a denervation of the basal ganglia with higher cortical regions of the SMN. This denervation or disconnection may result in decreased tolerance of the SMN towards consecutive perturbations. Decreased SMN NAT may in turn induce the clinical motor features of PD. Indeed, lower SMN and global NAT were associated with decreased motor performance independent of the level of DaT integrity. Overall, these findings expand on previous studies reporting dysfunctional connections to be primarily present between the dopamine-depleted putamen and the SMN ^43,44^. Further, the results overall support the notion that the dopamine-dependent disruption of functional connectivity between the putamen and the SMN primarily fuels motor impairments in PD, as previously suggested ^44^.

### Network attack tolerance as potential mechanism of motor reserve

Notably, in the current study we focused on general motor performance rather than the severity of singular clinical features, such as tremor, rigidity, or akinesia. This is particular important concerning our investigations in terms of NAT being a potential MR mechanism. MR is defined as the relative preservation of motor function despite progressive neurodegeneration, through the functional adaptation of large-scale neural networks ^2^. Here, we showed that independent of DaT loss, individuals with higher NAT in the global network and the SMN presented higher general motor performance. The current results expand on previous studies, which have mostly focused on regional structural ^45^ or functional ^4,5,46,47^ brain changes associated with MR. For example, greater functional connectivity within a network comprising the basal ganglia, the inferior frontal cortex, insula, and cerebellar vermis was associated with higher levels of MR (Chung et al., 2020). In addition, higher regional network integration in regions outside of the SMN has been suggested to be associated with MR ^4^. In contrast to these regional network measures, NAT describes network properties that are more representative of the brain’s global ability to compensate against neurodegeneration. Our findings highlight that the ability to preserve an efficient information flow within global and motor processing areas (i.e., SMN) despite DaT loss leads to a relative preservation of general motor performance. NAT may thus form the basis for underlying MR mechanisms and represent a quantitative measure of MR capacity in PD. Importantly, our findings even persisted when we corrected for clinical symptom severity. Thus, individuals with higher SMN NAT may be able to compensate for the effects of PD, independent of DaT loss and clinical severity.

Interestingly, in terms of cognitive function, we did not observe an effect of NAT in cognition-relevant networks on general cognitive performance. This may be due to the fact that we only included early PD patients with an absence of cognitive impairment. Hence, less variation in general cognitive performance may have led to the lack of findings concerning NAT and cognitive preservation. Nonetheless, a recent longitudinal study investigating NAT as a CR mechanism in two groups of PD patients with and without cognitive impairment reported that greater NAT of the fronto-parietal network served the maintenance of cognitive function over time ^7^.

The current findings thus suggest that NAT of distinct networks may serve as a resilience signature in PD relating to cognitive and motor reserve. Promoting these neural resources may provide means to slow disease progression. These neuronal resources may be promoted by means of interventional strategies such as trans-magnetic stimulation, but also through preventative measurements facilitating healthier lifestyles. Concerning the latter, lifestyle factors such as PA, education, occupation, or nutrition have consistently been associated with resilience capacity, predominantly in AD, but more recently also in PD ^1,45,48–50^. Yet, up to date, it remains unknown whether distinct lifestyle behaviors can promote NAT of certain neural networks in PD.

### The contribution of lifestyle factors to resilience and network attack tolerance

Aside from determining the association between NAT and DaT integrity and its role as potential resilience mechanism, we also aimed to examine the influence of two lifestyle factors, namely educational attainment, and lifestyle PA. Our analyses yielded that higher educational attainment was positively associated with NAT of the ATN. This is in line with previous findings indicating that higher levels of education are associated with greater functional connectivity of the dorsal attention network ^51^ and slower cognitive decline in healthy controls ^52^. Moreover, higher education was demonstrated to be linked to the build-up of neuronal resources delaying on one hand the onset of clinical symptoms in dementia ^53^ and on the other hand mitigating cognitive decline in PD ^12^. Interestingly, connectivity within the dorsal ATN appears to be crucial for cognitive performance. Lower connectivity levels of the dorsal ATN within the network itself and with the DMN and the FPN have been associated with the presence of mild cognitive impairment and decreased performance in attention and executive functions in PD ^54^. Thus, the build-up of greater NAT within the ATN due to higher levels of education may provide a compensatory mechanism to counteract cognitive decline in PD. Notably, we neither found an association between ATN NAT and general cognitive performance nor a moderating effect of education in our PD cohort. As outlined above, the lack of findings may be attributed to the minimal variation in general cognitive performance observed in a cohort of cognitively unimpaired PD patients. Future assessments, in more advanced PD patients may provide further insights into the role of educational attainment concerning the build-up of NAT and subsequent perseveration of cognitive performance.

Given that PD primarily is a movement disorder, we further considered another lifestyle factor that naturally appears closely related to motor function, namely lifetime PA. In a first step, we tested the direct effect of PA on the attack tolerance of distinct networks, which did not reach statistical significance. The lack of the direct effect of PA on NAT may potentially be caused by a more U-shaped association of PA with motor performance in PD. Indeed, it has been reported that higher lifetime PA delays the onset of PD but is linked to a more rapid decline in motor function from there on ^55^. Given the cross-sectional nature of our study, this assumption warrants further investigations in patients with early and advanced PD as well as longitudinal assessments. Nonetheless, in the current cohort of early PD patients, individuals with very high PA levels also demonstrated lower levels of motor performance pointing towards a more U-shaped association between lifestyle influences and disease progression.

Importantly, while we did not observe a direct effect of lifetime PA on NAT in any of the networks, we found a positive moderating effect of PA on the association between 1) SMN NAT and general motor performance, and 2) global NAT and general motor performance. Specifically, at similar NAT levels, PD patients with highly sedentary lifestyles portrayed worse general motor performance compared to PD patients with moderate PA levels. This suggests that a moderately active lifestyle, in contrast to a very sedentary one, may enhance the beneficial effects of NAT on motor performance. This is in line with other observations reporting a positive effect of premorbid PA on motor decline ^17^, and an association between dopamine depletion and motor impairment ^56^. One potential mechanism underlying this association could be the protective effect PA exerts on the basal ganglia’s structural integrity ^57^, and the beneficial effect of PA on the basal ganglia’s dopaminergic innervation ^18^. These effects may mitigate the severity of increasing striatocortical disconnection and thus aid the preservation of SMN NAT in the course of the disease ^5,44^. Notably, in our study this protective effect was not observed for higher PA levels. This observation may be attributed to the U-shaped association of PA with motor performance in PD discussed above. While several studies have already indicated that immediate, high-intensity exercise interventions positively impact PD symptomatology ^15,16,19,20^, our results further demonstrate that also lifetime, long-term PA, at least at moderate levels, contributes to the build-up of MR-associated mechanisms, such as global and SMN NAT, that mitigate the effects of the disease later in life.

Several limitations need be considered when interpreting the current results. First, despite this well-characterized cohort, the sample was relatively small and replication in a larger sample is warranted. Nonetheless, the tested models explained a relatively high amount of variance, which supports the validity of these results. Furthermore, we observed a significant effect of the three different fMRI sequences on global, SMN, and FPN NAT. Yet, this finding was likely driven by the imbalance in the number of sequences employed (5 vs. 16. vs. 1). Further, while GE is argued to be a superior measure of network integration, it may not capture longer, multiple paths that contribute to the integration of larger and more sparse networks since GE is based on the shortest path length ^58^. Hence, GE quantification of larger networks at lower network density levels (i.e., 5%), like in case of the tested global network (300 ROIs), may be less accurate. We aimed to circumvent this issue by assessing a wide range of densities. Finally, we focused on two reserve proxies, namely lifetime PA and education, which represent rather static and single measurements known to be associated with other lifestyle features such as socioeconomic status, occupation, lower chronic stress, and better general health. As more recently suggested ^59^, it may be advantageous to move away from investigating the role of single lifestyle factors to more cumulative measures including several lifestyle features when considering resilience mechanisms.

Notably, we additionally tested whether lifetime PA and educational attainment are correlated, which was not the case, suggesting an independent contribution of these factors to motor and cognitive reserve in PD. Moreover, in contrast to several other studies on MR in PD ^5,45^, we employed direct measures of resilience (PA and education) rather than using the indirect residual approach, which has recently been employed by several studies. The residual approach considers the variance in motor performance despite the level of dopaminergic deficit as proxy for reserve. While this approach may determine certain mechanistic principles explaining the clinical variance in PD patients, it does not allow to draw any conclusions about factors contributing to the observed variance ^2^. Therefore, it is of great importance to identify lifestyle features that promote resilience signatures, such as SMN NAT, which become relevant during the cause of a neurodegenerative disease. Identification of these factors may overall provide novel means in the prevention and intervention of these disorders.

### Conclusion

Taken together, the results of this study suggest that the brain’s compensatory capacity against neurodegenerative processes may be closely reflected by the degree of attack tolerance of distinct large-scale neural networks. Our findings underscore that lifelong behavioral factors, such as PA and education, affect the NAT of certain subnetworks, such as the somatomotor and attention network. Promoting these lifestyle factors may enhance the robustness of these networks, thereby contributing to the relative preservation of motor and cognitive performance despite the progressive loss of DaT in PD. Avoiding sedentary lifestyles and promoting physical activity may thus overall serve a low-cost interventional strategy to slow disease progression or delay the onset of PD.

## Author contributions

MH and AA were involved in outlining the research, planning the project, writing the manuscript, and making key decisions throughout the study. AA carried out all aspects of the data review, preprocessing and analysis, wrote the inhouse scripts utilized, carried out the statistical analysis, and created all figures and tables included. The data used was taken from the DoMoCo study. AA, MH, MB, VD, HT and TvE were highly involved in the execution of this study (e.g., data acquisition, patient recruitment, study design, data management). NH and KM contributed by guiding and overseeing the statistical approach. EG established the analytical framework for calculating the lifetime physical activity score. Finally, all authors reviewed the final manuscript

## Acknowledgements

This work was supported by the Deutsche Forschungsgemeinschaft (DFG, German Research Foundation) – Project-ID 431549029 – SFB 1451/C03. HT was supported by the Cologne Clinician Scientist Program (CCSP)/ Faculty of Medicine/ University of Cologne, funded by the DFG – Project ID 413543196. Furthermore, we are grateful for the patience and commitment of every participant enrolled and contributing to the DoMoCo study. We would like to thank the Human Motor Assessment Center (Project Z03 of the SFB 1451) for providing data.

## Data availability

The data supporting this study’s findings are findable in the CRC1451 data registry (https://www.crc1451.uni-koeln.de/), and reasonable requests can be addressed to the corresponding author.

