## SupplementaryMaterial for "Lifetime physical activity and network attack tolerance contribute to the preservation of motor function in Parkinson’s disease"

**Supplementary Material:**

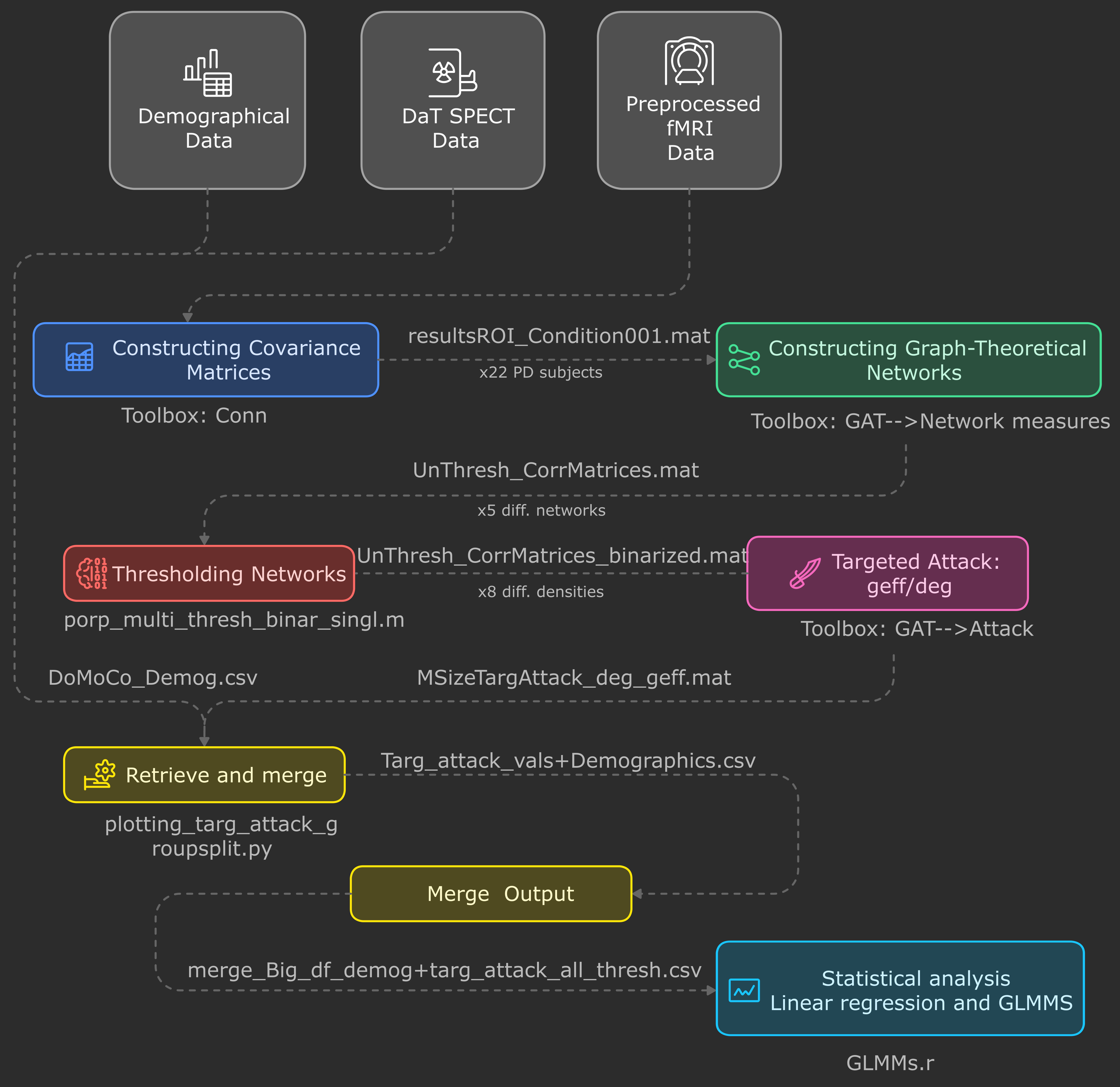

**S-Fig 1.: Procedural map of the analysis pipeline implemented**. Grey boxes represent the data that served as an input to the pipeline. The colored boxes represent each step undertaken throughout the analysis. Underneath each box, the toolbox or script used for this step is depicted. The text above the arrows indicates the data format generated as an output and subsequently served as an input in the next step. If this step was carried out multiple times, it is mentioned below the arrow.

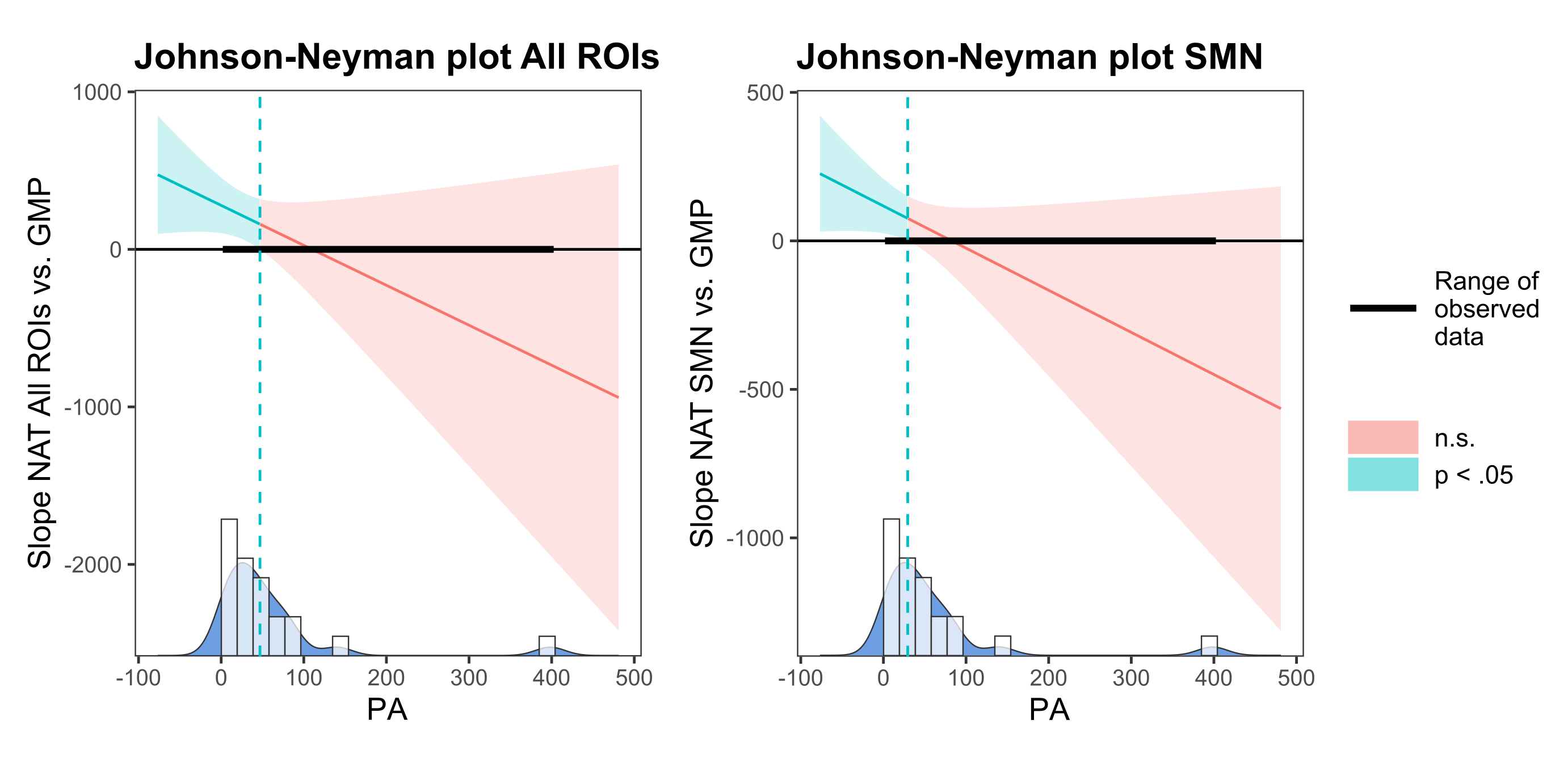
**S-Fig 2.: Johnson-Neyman plots** assessing the moderating effect of PA on the association between general motor performance (GMP) and global network attack tolerance (NAT) (left) and the association between GMP and somatomotor network (SMN) NAT (right). Ranges where PA significantly moderated the effect on the association between NAT and GMP are shown in light blue. Ranges of non-significance are depicted in red. The bold black bar indicates the range of observed data. On the x axis a ‘densigram’ (density in dark blue and histogram in white) shows the distribution of observed PA levels, providing context for the applicability of the identified ranges of significance. Significance threshold: p < .05.

**Link to Pictograms used:**

Cognitive performance: <https://www.freepik.com/icon/cognitive-function_11604264>

Motor function: <https://iconscout.com/icon/fine-motor-skills-5149837>
